# Immuno-genomic Pan-cancer Landscape Reveals Diverse Immune Escape Mechanisms and Immuno-Editing Histories

**DOI:** 10.1101/285338

**Authors:** Shinichi Mizuno, Rui Yamaguchi, Takanori Hasegawa, Shuto Hayashi, Masashi Fujita, Fan Zhang, Youngil Koh, Su-Yeon Lee, Sung-Soo Yoon, Eigo Shimizu, Mitsuhiro Komura, Akihiro Fujimoto, Momoko Nagai, Mamoru Kato, Han Liang, Satoru Miyano, Zemin Zhang, Hidewaki Nakagawa, Seiya Imoto, on behalf of the PCAWG Mitochondrial Genome and Immunogenomics Working Group and The PCAWG Network

## Abstract

Immune reactions in the tumor micro-environment are one of the cancer hallmarks and emerging immune therapies have been proven effective in many types of cancer. To investigate cancer genome-immune interactions and the role of immuno-editing or immune escape mechanisms in cancer development, we analyzed 2,834 whole genomes and RNA-seq datasets across 31 distinct tumor types from the Pan-Cancer Analysis of Whole Genomes (PCAWG) project with respect to key immunogenomic aspects. We show that selective copy number changes in immune-related genes could contribute to immune escape. Furthermore, we developed an index of the immuno-editing history of each tumor sample based on the information of mutations in exonic regions and pseudogenes. Our immuno-genomic analyses of pan-cancer analyses have the potential to identify a subset of tumors with immunogenicity and diverse background or intrinsic pathways associated with their immune status and immuno-editing history.

## Introduction

Genomic instability and inflammation or immune responses in the tumor microenvironment are major underlying hallmarks of cancer (Hanahan and Weinberg, 2011), and the interaction between the cancer genome and immune reactions could have important implications for the early and late phases of cancer development. The immune system is a large source of genetic diversity in humans and tumors (Lefranc et al., 1999). Human leukocyte antigen (HLA), the vast number of unique T‐ and B-cell receptor genes, and somatic alterations in tumor cell genomes enable the differentiation between self and non-self (tumor) via neoantigen (NAG) presentation, which contributes to positive or negative immune reactions related to cancer (Linnemann et al., 2015; Tran et al., 2014; Kreiter et al., 2015; Yarchoan et al., 2017). A variety of immune cells can infiltrate tumor tissues and suppress or promote tumor growth and expansion after the initial oncogenic process (Grivennikov et al., 2010). Such cancer immuno-editing processes (Schreiber et al., 2011) sculpt the tumor genome via the detection and elimination of tumor cells in the early phase and are also related to the phenotype and biology of developed cancer. It is important to investigate escape mechanism of tumor cells from immuo-editing and also whether microenvironment around tumors help the escape or not. If the microenvironment helps it, it is largely unclear how the immune microenvironment helps tumor cells with or without genetic alterations of immune molecules escape immuno-editing, and methods to observe the immuno-editing history in clinical human tumors are needed.

Emerging therapies targeting immune checkpoint or immune-escape molecules are effective against several types of advanced cancer (Sharma et al., 2011; Pardoll, 2012; Mahoney et al., 2015; Zaretsky et al., 2016; Anagnostou et al., 2017); however, most cancers are still resistant to these immunotherapies. Even after successful treatment, tumors often acquire resistance via another immune escape mechanism or by acquiring genomic mutations in intrinsic immuno-signaling pathways, such as the IFN gamma pathway or MHC (HLA) presentation pathway, related to NAG (Gao et al., 2016; Shin et al., 2017). Tumor aneuploidy is also correlated with immune escape and the response to immunotherapy (Dovoli et al., 2017); hence, for comprehensively understanding of cancer immunology and its diversity, whole genome analysis is necessary. We here analyzed the whole genome sequencing (WGS) of 2,834 donors and RNA-seq data from Pan-Cancer Analysis of Whole Genomes (PCAWG) project (Campbell et al., PCAWG marker paper) with respect to key immunogenomic aspects using computational approaches (Hackl et al., 2016). Our results demonstrate that diverse genomic alterations in specific tumor types, variation in immune microenvironments, and variation in oncogenic pathways are related to immune escape, and we further observed immune-editing during cancer development. To illustrate the history of immunoediting history for each cancer genome and to explore underlying molecular pathways involved, we defined immuno-editing indexes (IEIs) by comparing exonic NAGs to virtual NAGs in pseudogenes.

## Results and Discussion

Based on recent intensive investigations of the relationship between copy number alterations (CNAs) and cancer development and progression (Dovoli et al., 2017, Laumont et al., 2018), CNAs may associate with immunological profiles of cancers, however causality is largely unknown. Somatic alterations in immune-related genes may contribute to cancer development and progression or immune escape in certain solid tumors and hematopoietic tumors. To investigate the effect of genomic alterations occurred in immune system, we compiled a list of 267 immune-related genes (Supplementary Table 1) that could be assigned to four categories: the immune escape pathway, antigen presentation pathways for *HLA* class I and *HLA* class II, and the cytokine signaling and apoptotic pathways, including genes involved in the IFN gamma pathway. An analysis of PCAWG consensus variant calls (Campbell et al., PCAWG marker paper) demonstrated that most tumor samples have at least one somatic alteration in these immune-related genes (Figure 1a). Although CNAs were the most frequently detected type of somatic alteration, many point mutations and structural variants (SVs) were also detected in the immune-related genes including *HLA-A, HLA-B, HLA-C* and *B2M*, Beta-2 microglobulin (Supplementary Figure 1a).

**Supplementary Table 1:**
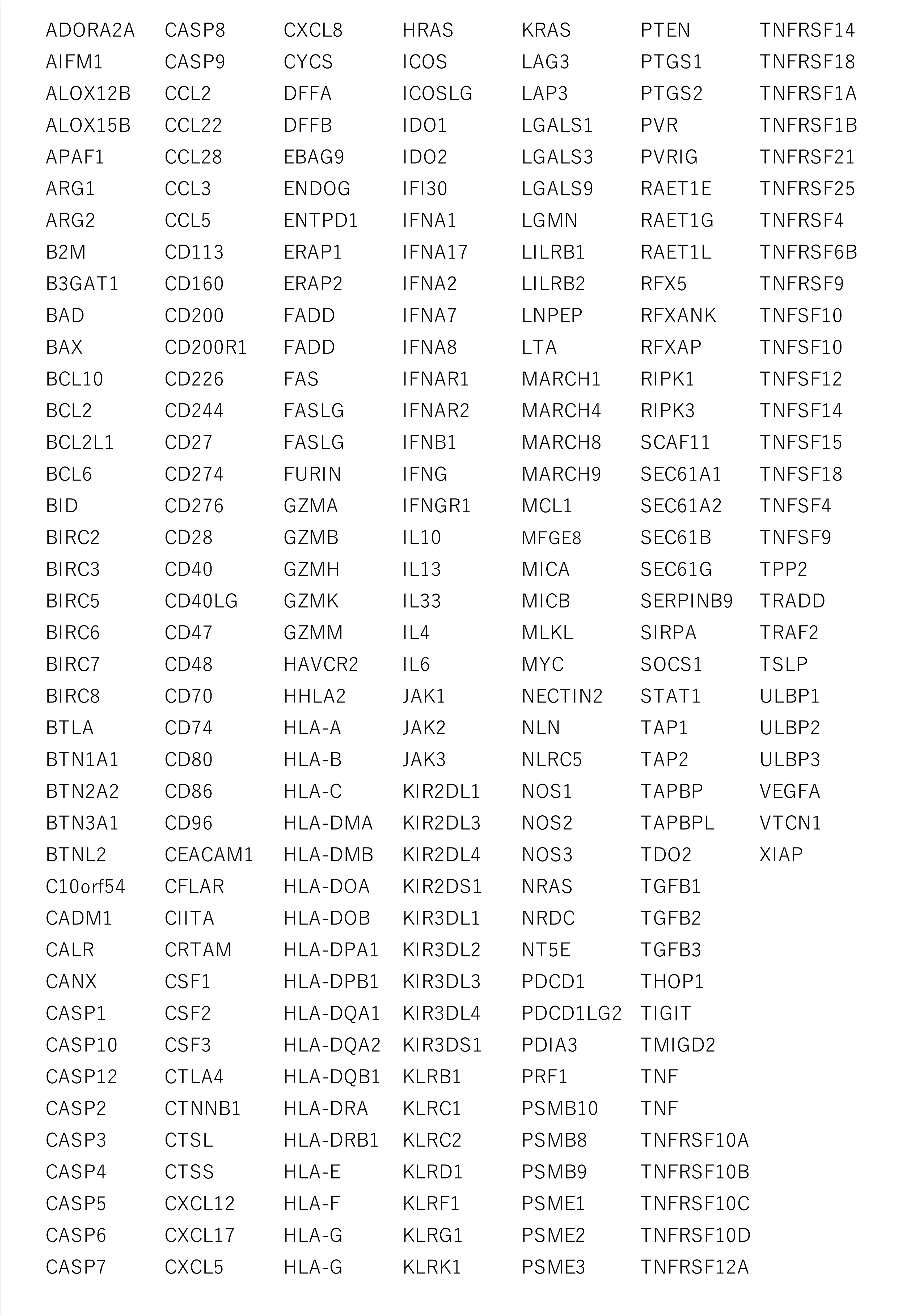
List of analyzed immune-related genes.

**Figure 1:**
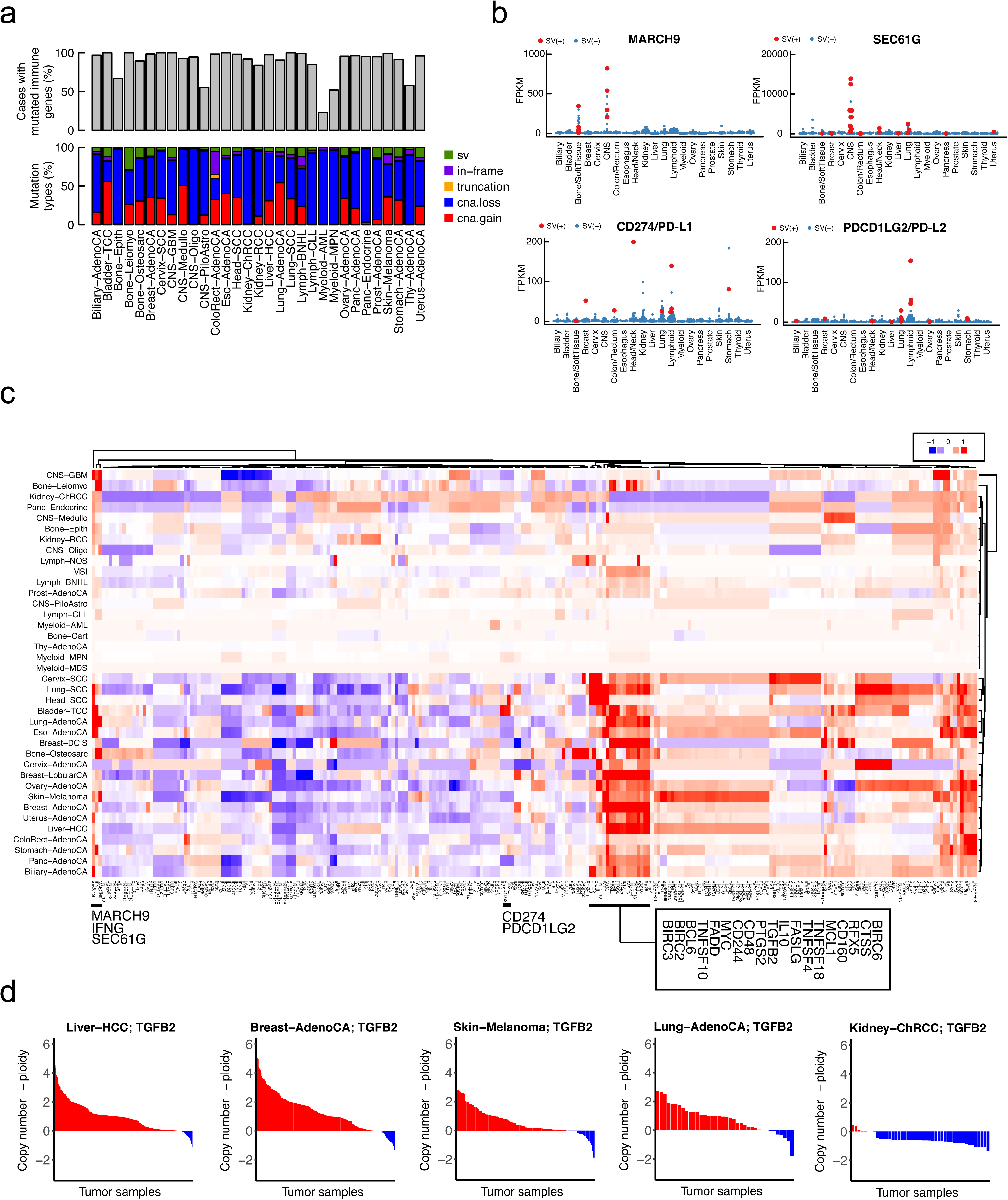
Mutation landscape of immune-related genes. (a) Frequency and types of somatic mutations in immune genes. Single nucleotide variants (SNVs), insertions and deletions (indels), structural variants (SVs), and copy number alterations (CNAs) were examined in immune genes and donors for multiple types of tumors, where ‘truncation’ represents ‘stop gain SNV’ and ‘frameshift indel,’ and ‘in-frame’ means ‘nonsynonymous SNV’ and ‘in frame indel.’ (b) Overexpression of immune-related genes and its association with SVs in each tumor type. Red and blue dots represent tumor samples with and without SVs, respectively. (c) Copy number of *IL10* offset by tumor ploidy. Tumor samples are colored red and blue to indicate whether the copy number is above or below the ploidy level, respectively. (d) Copy number of *IL10* offset by tumor ploidy. Tumor samples are colored red and blue to indicate whether the copy number is above or below the ploidy level, respectively.

**Supplementary Figure 1:**
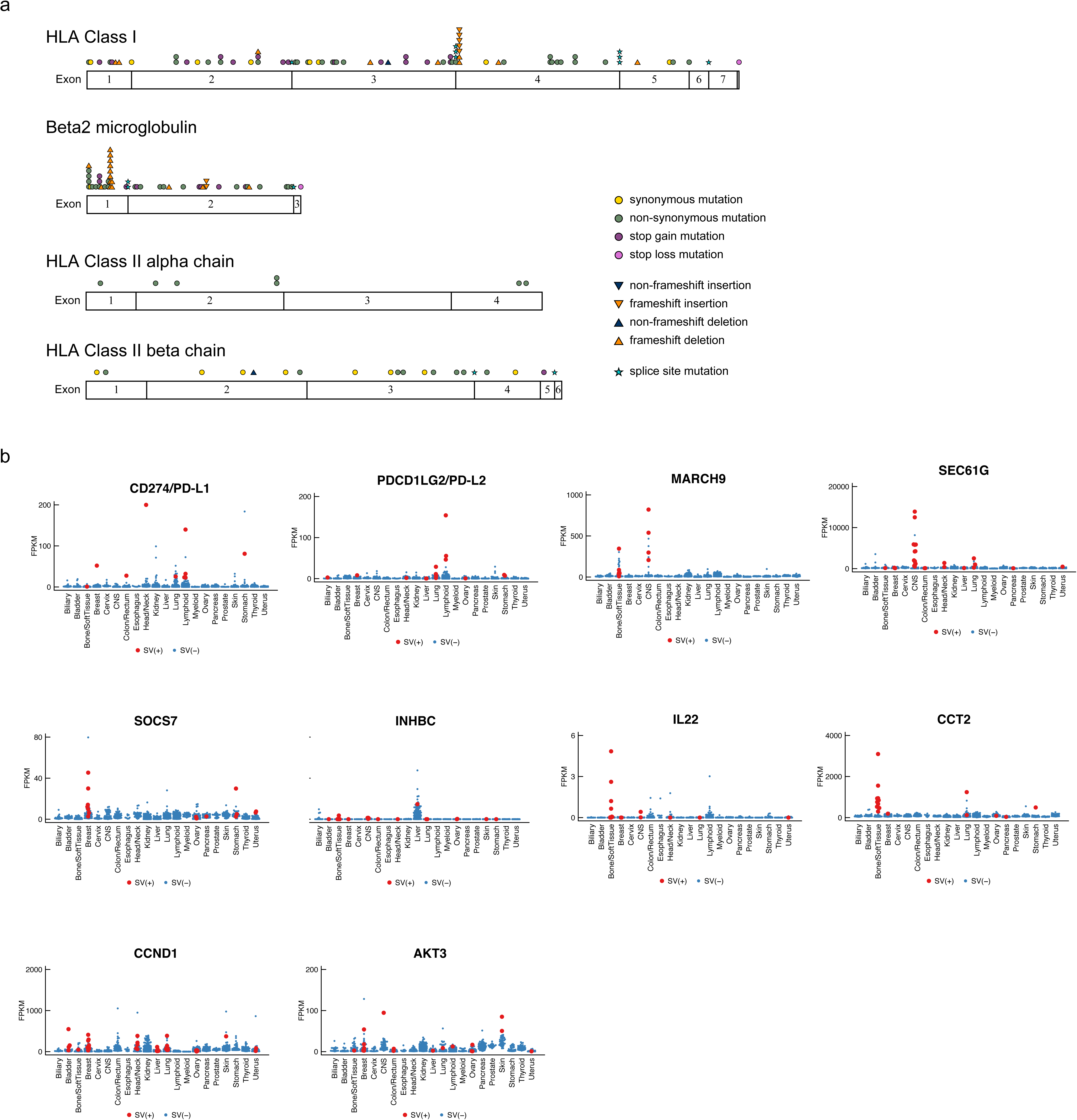
Identified somatic mutations in HLA and B2M genes (a) and structural variant-related overexpression of ten immune-related genes (b).

We also investigated SVs (Zhang et al., PCAWG marker paper) in immune-related genes. Although SVs are relatively rare compared to CNAs, they may have a large impact on the expression and function of affected genes, as exemplified by a recent report of the 3?-untranslated region of *CD274/PD-L1* (Kataoka et al., 2016). For each immune-related gene, we compared mRNA expression levels between SV-positive and SV-negative cases. For ten immune-related genes (*CD274/PD-L1*, *PDCD1LG2/PD-L2*, *MARCH9*, *IL22*, *SEC61G*, *CCND1*, *CCT2*, *INHBC*, *AKT3*, and *SOCS7*), we detected a statistically significant association between the occurrence of SVs and the upregulation of expression (q-value < 0.05; Figure 1b and Supplementary Figure 1b). *PDCD1LG2*/*PD-L2* can interact with PD-1 and PD-L1, resulting in inhibitory signals that modulate the magnitude of T-cell responses (Latchman et al., 2001; Rozali et al., 2012). *MARCH9*, an E3 ubiquitin ligase, downregulates MHC class II molecules in the plasma membrane (Janke et al., 2012), and *SEC61G* is involved in the translocation of HLA class I proteins to the endoplasmic reticulum for clearance (Albring et al., 2004). These findings indicate that SVs could affect HLA complexes and their expression or activity/clearance as well as immune checkpoint molecules, which may facilitate the immune escape of tumor cells.

As shown in Figure 1a, CNAs are the most frequently observed alterations in immune-related genes. Cancers harboring many CNAs tended to show less immune involvement and worse responses to immunotherapies (Davoli et al., 2017), and this can potentially be explained by CNAs in immune-related genes. We next compared the copy numbers of immune-related genes with the ploidy levels of tumors to differentiate between selective increases in copy number or changes in ploidy or averaged changes of chromosomes.

We first analyzed other immune-related genes and tumor types, including MSI (microsatellite instability)-positive tumors (Fujimoto, PCAWG-7, et al., BIORXIV/2018/406975) with strong immunogenicity (Le et al., 2015) due to high numbers of NAGs. For each immune-related gene, we used *t*-tests to evaluate whether the copy number differences from the ploidy level are significant or not in each tumor type. The results are summarized as a landscape of selective copy number changes in Figure 1c (showing the mean copy number changes against the ploidy value) and Supplementary Figure 2 (showing the statistical significance of selective copy number changes).

We focused on one distinctive cluster in Figure 1c, which seems to be occurrence of selective copy number gain in multiple tumor types; transforming growth factor beta 2 (*TGFB2*) and interleukin-10 (*IL10*) were included in this cluster. *TGFB2* and *IL10* are located on chromosome *1q* and both function as suppressors of immune cells (Itakura et al., 2011; Wiguna and Walden, 2015; Yang et al., 2015). *IL10* expresses not only immune cells, but also tumors; the functions of IL10 produced from tumor cells were mainly reported in melanoma. The selective copy number gains for these immune immune genes are likely to be related to tumor-immune system interactions (Figure 1c). Recently, a molecule simultaneously inhibiting *TGFB2* and *PD-L1* is reported and showed its high efficacy (Lan et al., 2018; Strauss et al., 2018).

We then examined the differences between copy number of *TGFB2* and ploidy level, relative copy number of TGFB2, for each donor of multiple tumor types (Figure 1d). In Liver-HCC, Breast-AdenoCA, Skin-Melanoma, and Lung-AdenoCA samples, the *TGFB2* copy number was specifically increased, rather than the ploidy level, in almost all tumors. Since *TGFB2* functions as a repressor of immune cells, the amplification or gain of *TGFB2* is possibly related, in part, to the immune escape mechanism. However, in Kidney-ChRCC, no significant selective amplification was observed.

It might be important to know an immune escape mechanism related to *TGFB2*. *SEC61G* and *MARCH9*, both of which exhibited significant overexpression related to SVs (Figure 1b), showed different patterns from those of *TGFB2* and *IL10*. *MARCH9* showed statistical significance in some tumor types; considering the mean value of the differences in each tumor type, selective copy number gain was detected in CNS-GBM and Bone-Leiomyo. Additionally, *SEC61G* was selectively amplified in CNS-GBM and Head-SCC. Interestingly, donors with SV-related overexpression and donors with selective copy number gains were highly correlated; however, selective copy number gain could only partially explain the overexpression of these genes for the donors without SVs (Supplementary Figure 3).

**Supplementary Figure 2:**
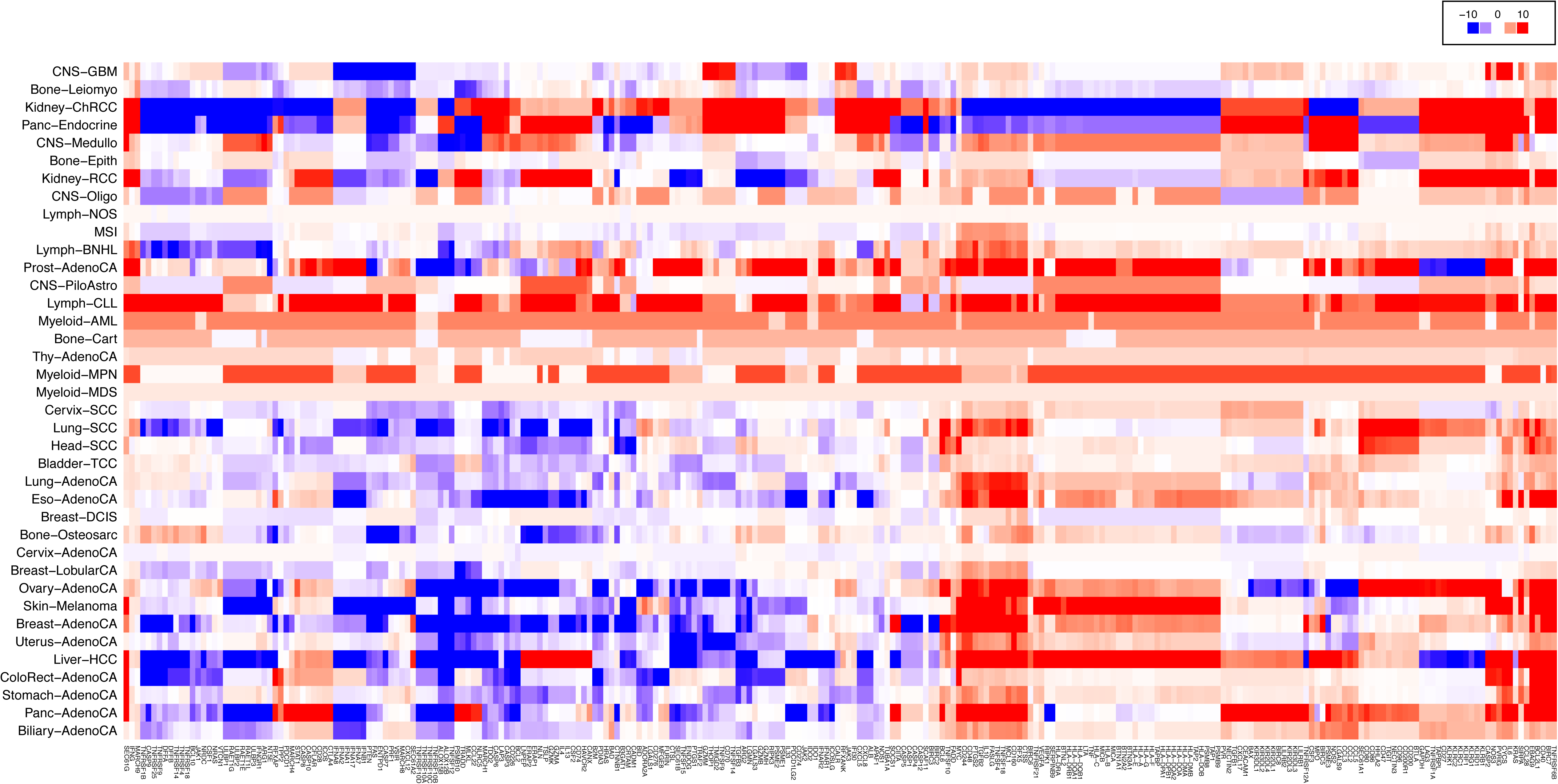
Statistical significance of selective copy number changes. The color of each element represents the score of the statistical test, defined by ‐sign(*t*-statistic)*log10(p-value). The function sign(a) takes +1 if a is positive, otherwise ‐1.

**Supplementary Figure 3:**
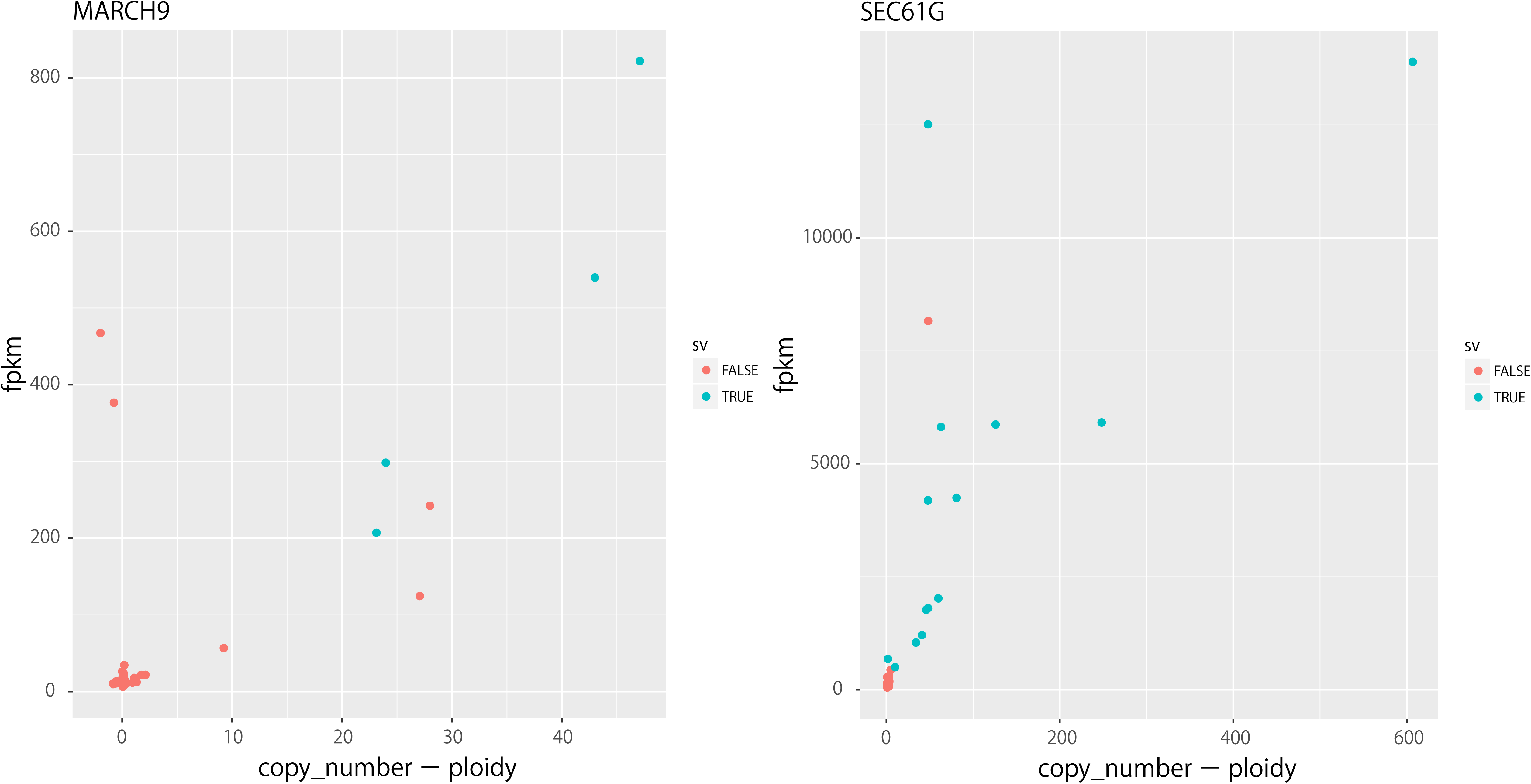
Selective copy number gain and structural variation can explain RNA overexpression.

In Skin-Melanoma, the copy numbers of genes on chromosome 6, including HLAs, were significantly greater than the ploidy level (p = 2.26E-10 for *HLA-A*), which could paradoxically increase immune pressure. However, the copy number of *IL10* was also significantly (p = 8.1E-10) and selectively increased, potentially contributing to escape from immune pressure. By contrast, in Kidney-ChRCC and Panc-Endocrine samples, the copy numbers of *HLAs* compared with the ploidy level show the opposite tendency, and *IL10* follows this. Since *HLAs* are not selectively increased, the copy number gain for *IL10* may be unnecessary for immune escape. In Lymph-NOS and Myeloid-MDS, copy numbers of almost all immune genes were consistent with ploidy and were not selectively changed (minimum p = 0.498 and 0.184 for Lymph-NOS and Myeloid-MDS, respectively). MSI tumors showed weak selective copy number increases for genes in the cluster including *IL10* (p = 0.000644); however, significant results were not obtained for other immune-related genes. In these tumor types, there may exist different immune escape systems, other than the selective copy number gain of these immune genes.

We analyzed the differences between copy number and ploidy and, interestingly, found that genomic regions containing genes that function as suppressors of the immune system, such as *TGFB2* and *IL10,* are selectively increased in many types of tumors. Copy number gains of these immune-related genes could arise and be selected during the establishment of immune escape. Therefore, selective copy number gains may be involved in the history of immune escape. Since *TGFB2* and *IL10* could play important roles in immune escape based on their function, our findings indicate that selective copy number gain is a remarkable system in the mechanism of immune escape. However, no selective copy number gains were observed in immune checkpoint genes, *i.e.*, *PD-L1* and *PD-L2*, which function as part of the immune escape mechanism, further suggesting the diversity of immune escape mechanisms.

During tumorigenesis, mutant peptides derived from nonsynonymous somatic mutations are presented by HLA molecules to T cells (Figure 2a) (Robbins et al., 2013; Carreno et al., 2015). Although these NAGs serve to eliminate tumor cells, some cells escape this immune surveillance and eventually contribute to the formation of clinical tumors (Figure 2b) (Burnet, 1970; Dunn et al., 2002). To estimate the strength of immune surveillance or immune pressure experienced by tumor cells in each sample, we developed a novel approach to measure the strength of immune pressure using pseudogenes as an internal control of each tumor (Figure 2a) (see Methods); those are not translatable. First, we identified predicted NAGs from somatic substitutions in exonic regions of whole genome sequences and compared them to those similarly derived from pseudogenes (Supplementary Figure 4). In this process, we used the HLA types (class I and II, shown in Supplementary Figure 5) determined by our new pipeline, referred to as ALPHLARD (see Methods). The accumulation of somatic mutations in exonic regions versus virtual somatic mutations in pseudogenes during tumorigenesis is schematically represented in Figure 2b. If tumor cells grew under strong immune pressure, the difference between predicted NAGs in exonic and pseudogene regions would be large. This difference is expected to be small if tumor cells immediately escape from immune pressure in the carcinogenic process (Figure 2c). We defined the immuno-editing index (IEI) according to this concept (see Methods). The virtual neoantigen ratio R*_P_* for mutations in pseudogene regions and the neoantigen ratio R*_E_* for exonic regions can be plotted (Figure 2d) to determine the immune pressure for each tumor sample. IEI is defined as the log-ratio of R*_P_* to R*_E_*. We used IEI to characterize the histories of different donors, including immuno-edited and immuno-editing-resistant tumors.

**Figure 2:**
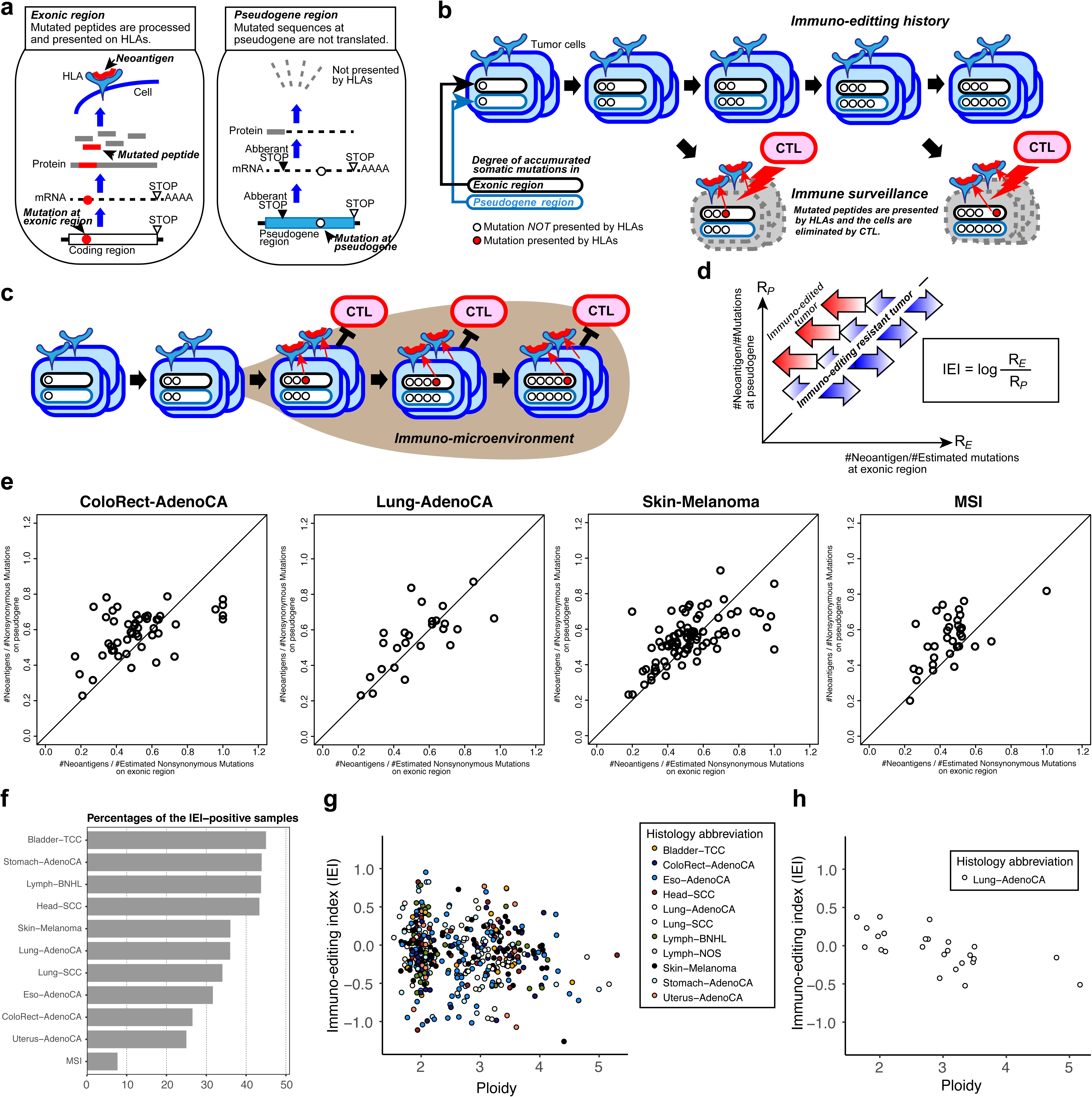
Analysis of immuno-editing history. (a) Overview of the presentation of neoantigens generated from nonsynonymous mutations in exonic regions. Pseudogene regions are not translated and mutations that accumulate in pseudogenes are not presented by the HLA complex. (b) Relationship between accumulated mutations in exonic regions and pseudogenes in the immunoediting history. Although CTLs (cytotoxic T-cells) eliminate tumor cells by recognizing these NAGs, some tumor cells escape this immune surveillance mechanism and eventually contribute to the formation of a clinical tumor. (c) In immuno-editing-resistant tumors, the tumor cells immediately escaped from immune pressure in the carcinogenic process, and the difference between NAGs in exonic and pseudogene regions was expected to be small. (d) Immuno-pressure plot of (virtual) neoantigens in exonic regions and psueodogenes. The *x*-axis represents the virtual neoantigen ratio R*_P_* for mutations in pseudogene regions and the *y*-axis shows the neoantigen ratio R*_E_* in exonic regions. IEI (immuno-editing index) was defined as the log ratio of R*_P_* to R*_E_* and was used to characterize the immune-editing history of each donor, with immuno-edited and immuno-editing-resistant tumors. (e) Immuno-pressure plots of four cancer types. MSI-positive tumors show the most immuno-edited tumor characteristics; in other cancers, many tumors showed an immuno-editing-resistant tendency. (f) The proportion of immuno-editing-resistant tumors. (g, h) Tumor ploidy and IEI for a pan-cancer analysis (g; n = 433) and lung adenocarcinoma (h; n = 25). Each dot represents a tumor sample.

**Supplementary Figure 4:**
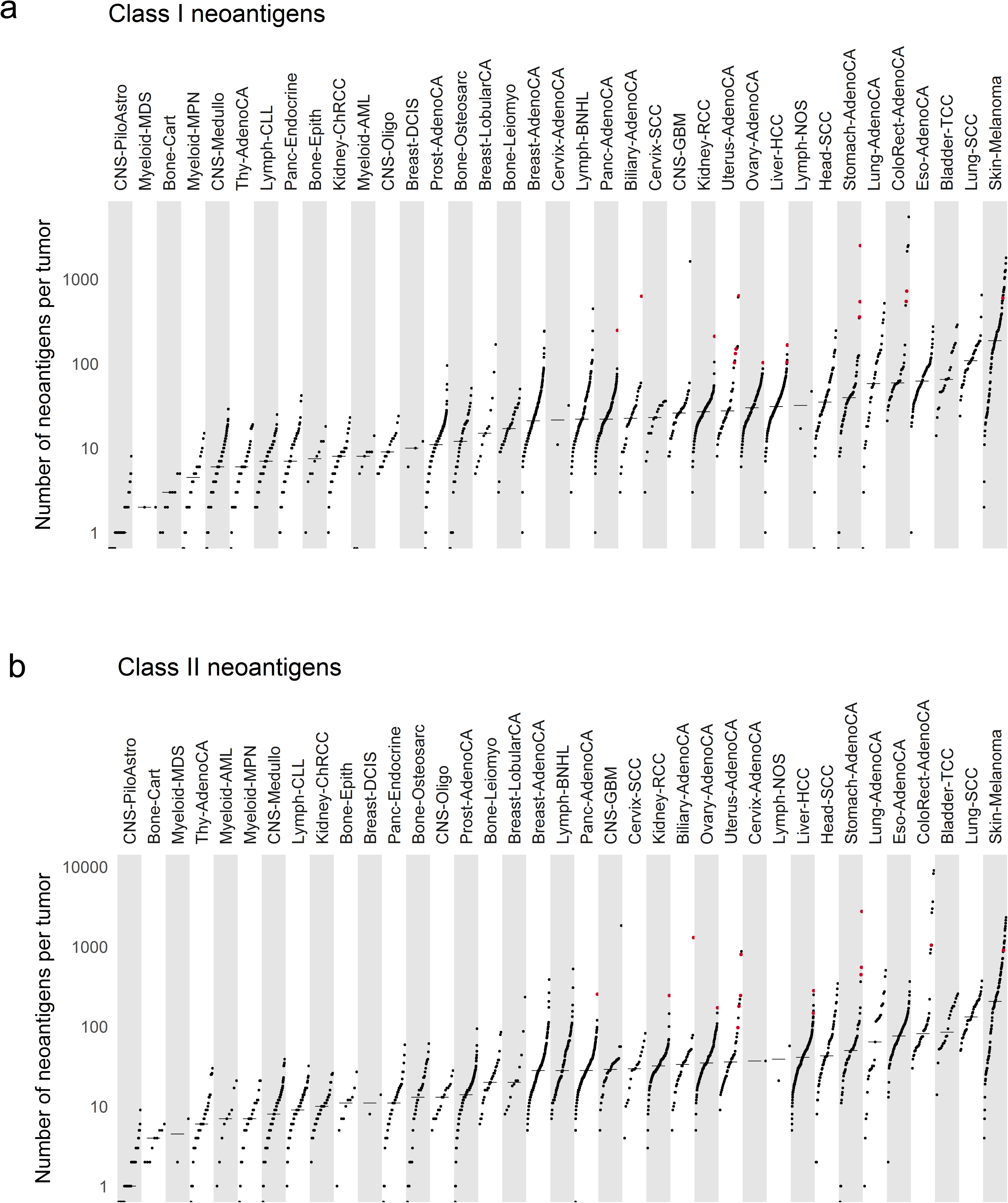
Distributions of neoantigens for each tumor type. (a) Class I and (b) class II.

**Supplementary Figure 5:**
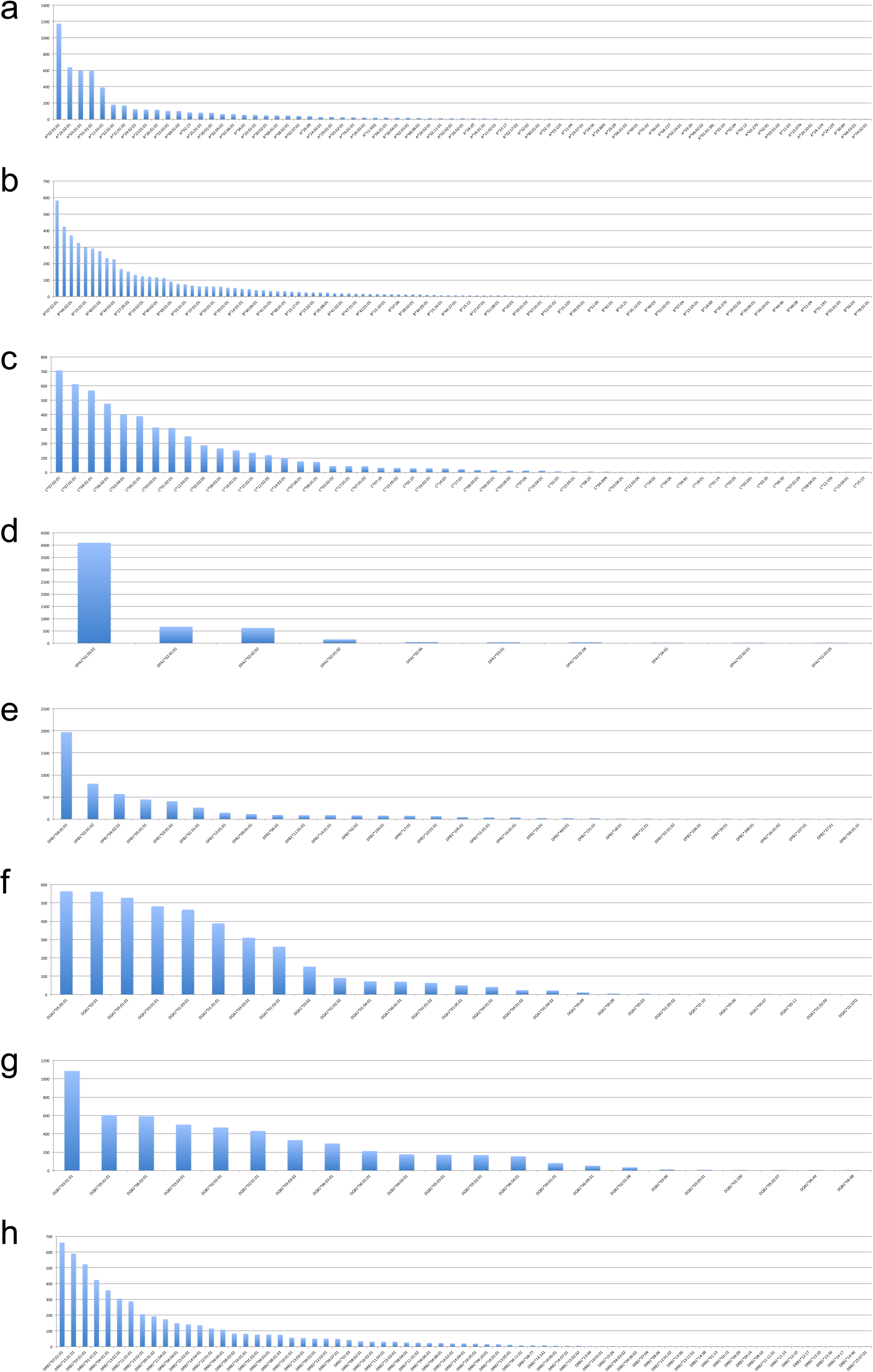
Distributions of determined HLA types from whole genome sequence data.

In subsequent analyses, we investigated the history of immune pressures for multiple tumor types, as revealed by IEI. The distributions of immune pressure for four cancers are shown in Figure 2e. The percentage of IEI-positive samples, *i.e.*, immune-editing-resistant tumors, in each tumor is shown in Figure 2f. MSI-positive tumors show immuno-edited tumor characteristics, suggesting that MSI-positive tumors were continuously being under strong negative selection from the immune system. Bladder-TCC, Stomach-AdenoCA, Lymph-BNHL and Head-SCC samples showed immuno-editing-resistant tendencies, indicating that mutations generating NAGs were removed by negative selection during tumorigenesis.

We compared the IEI values with the ploidies using pan-cancer data and observed a significant negative correlation (Pearson’s correlation coefficient, *r* = −0.13, p = 0.0051) (Figure 2g). Among the 11 tumor types, the strongest correlation was observed in Lung-AdenoCA (*r* = −0.66, p = 0.00028), and multiple tumor types, including ColoRect-AdenoCA, Eso-AdenoCA, and Skin-Melanoma, showed weak negative correlations, although these were not statistically significant. The negative correlation between IEI and ploidy can likely be attributed to the scenario in which a copy number gain leads to high expression of NAGs and thus high immune pressure.

We next examined the immune characteristics or signatures related to the difference in immune escape histories (as determined by IEI). Differentially expressed genes between IEI-positive and - negative tumors were analyzed to find acquired phenotypes or micro-environmental characteristics that promote tumor cell escape from immune pressure. Using four signatures related to immune characteristics, *i.e.*, HLA class I, cytotoxic, immune checkpoint, and cell component, we divided samples into two groups, referred to as hot (inflamed; presence of infiltrating immune cells) and cold (non-inflamed) tumors, and we further used gene set enrichment analyses (GSEA) to elucidate the pathways (Figure 3a). The genes related to each of above four signatures are listed in Supplementary Table 2. Interestingly, we found distinct patterns of gene set enrichment in hot and cold tumors (Supplementary Figure 6). In the hot tumors, multiple gene sets, *e.g.*, interferon gamma response and inflammatory response genes, were commonly enriched in most tumor types (Supplementary Figure 6). In contrast, in the cold tumors, most gene sets were differentially enriched in a tumor-specific manner. Thus, immune escape pathways preventing immune-cell infiltration, *i.e.*, cold tumors, could be diverse and specific to each tumor type. Recently, Eynden et al. (2018) discussed the neoantigen depletion in various tumors. A difference of their work and ours is that we focused on the strengthen of immune pressure for individual tumors, whereas they analyzed characteristics of each tumor type. They concluded that the signal of negative selection is not strong or absent in most tumor types, however interestingly they found that only lung adenocarcinoma showed significant negative selection, which is consistent to our result.

**Figure 3:**
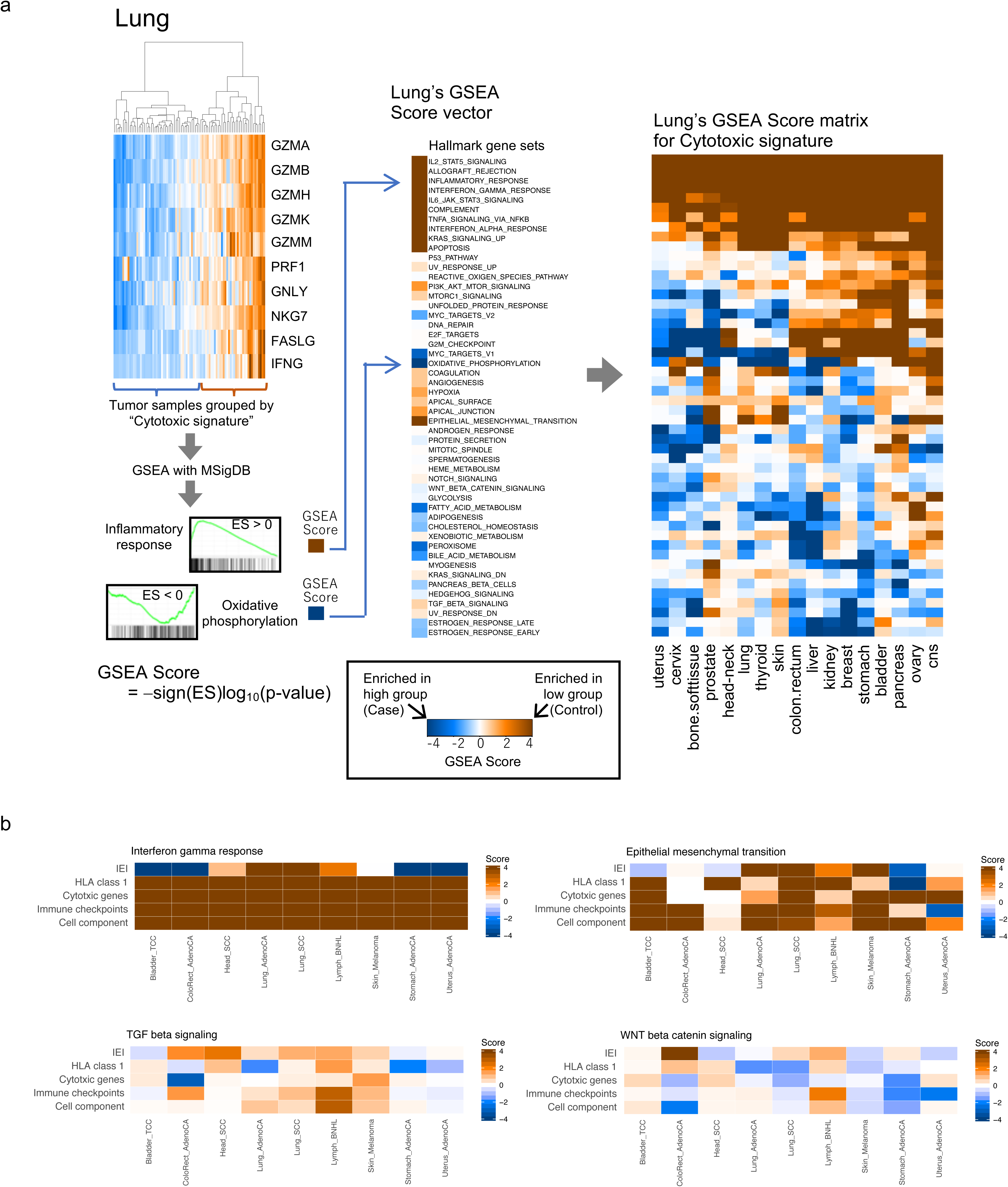
Analysis of immune signatures. (a) The samples are clustered based on the expression of genes listed in Supplementary Table 2 and the expression of genes in two groups of samples were compared using two-sided *t*-tests. The enrichment of gene sets defined by MSigDB was evaluated by GSEA. The score is defined in the same way as selective copy number changes, using the sign of the enrichment score and its p-value. (b) GSEA in four gene sets (‘interferon gamma response,’ ‘EMT,’ TGF-beta signaling,’ and ‘WNT/β-catenin signaling’) was used to determine the degree of enrichment of the four immune signatures and IEI. The color of each pair of tumor type and gene set represents the GSEA score (Figure 3a)

**Supplementary Figure 6:**
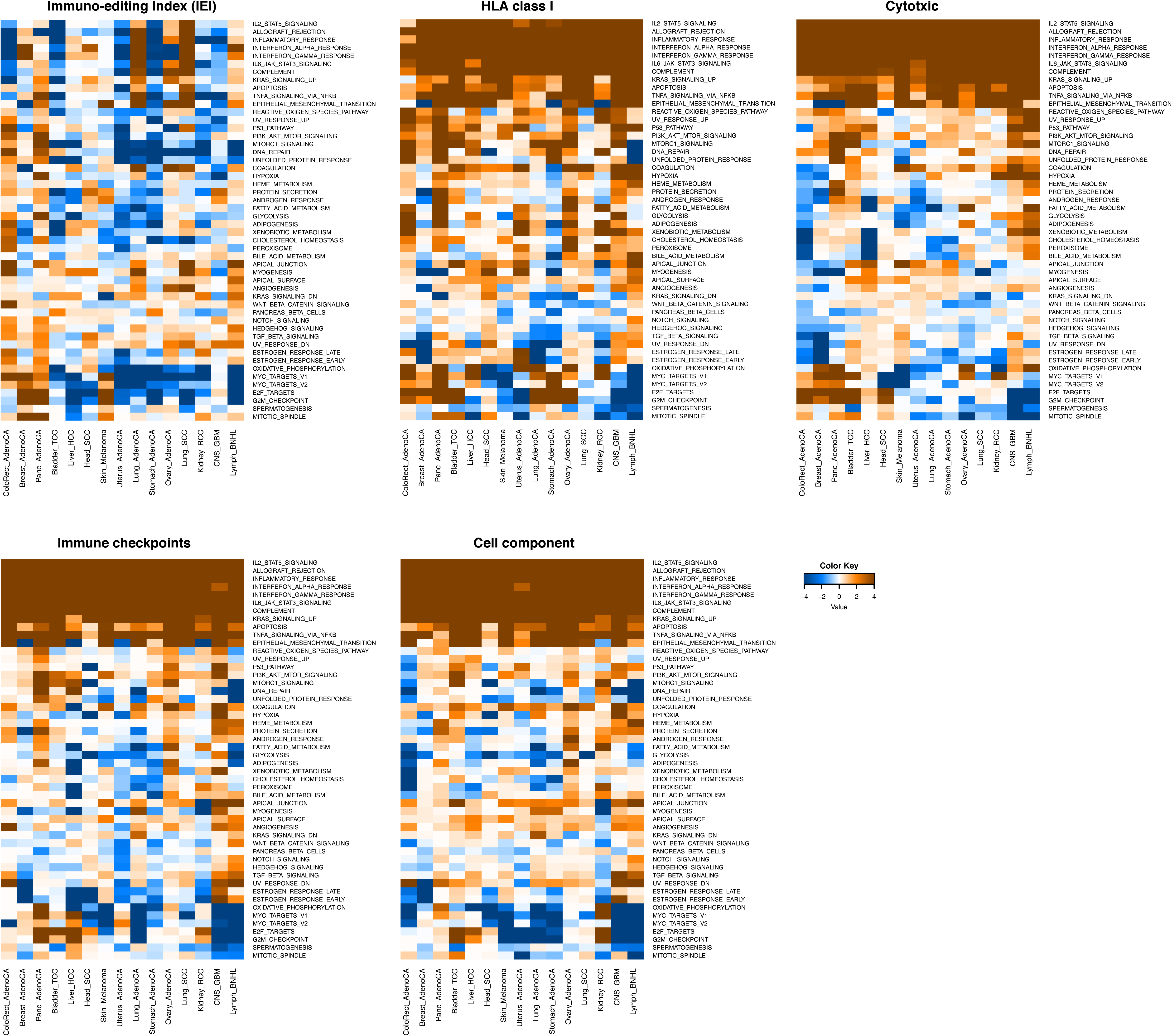
For IEI (positive and negative) and four immune signatures for tumor immuno-types (hot and cold), GSEA results for all gene sets and tumor types are summarized.

**Supplementary Table 2:** Genes involved in four immune signatures.

| HLA class I | Cytotoxic | Immune checkpoints | Cell component |
| --- | --- | --- | --- |
| HLA-A | GZMA | PDCD1 | PTPRC |
| HLA-B | GZMB | CTLA4 | CD2 |
| HLA-C | GZMH | HAVCR2 | CD3G |
| B2M | GZMK | LAG3 | CD3E |
| PSMB8 | GZMM | BTLA | CD4 |
| PSMB9 | PRF1 | TIGIT | CD5 |
| PSMB10 | GNLY | CD96 | CD7 |
| TAP1 | NKG7 | CD200R1 | CD8A |
| TAP2 | FASLG | LILRB1 | KLRK1 |
| TAPBP | IFNG | LILRB2 | KLRB1 |
| NLRC5 |  | CD160 | KLRD1 |
|  |  |  | CD19 |
|  |  |  | MS4A1 |
|  |  |  | ITGAX |
|  |  |  | ITGAM |
|  |  |  | CD14 |
|  |  |  | CD33 |

Furthermore, we specifically examined the degree of enrichment of four specific gene sets with respect to IEI (for gene sets of interferon gamma response, EMT (epithelial to mesenchymal transition) (Terry et al., 2017), TGF beta signaling (Yang et al., 2015), and WNT/β-catenin signaling (Spranger et al., 2014; Pai et al., 2017) (Figure 3b). The interferon gamma response gene set was enriched in inflamed tumors of all tumor types, as expected, as the expression levels of these genes are higher in inflamed tumors than in non-inflamed tumors. Using IEI, this trend was maintained in Head-SCC, Lung-AdenoCA, Lung-SCC, and Lymph-BNHL samples; in these tumor types, those genes are more highly expressed in the IEI-positive group than in the IEI-negative group. However, in Skin-Melanoma, the enrichment of this gene set was not significant, while these genes were significantly underrepresented in four tumor types (Bladder-TCC, ColoRect-AdenoCA, Stomach-AdenoCA, and Uterus-AdenoCA). This suggests that the diversity of immuno-editing histories depends on the tumor type. For the EMT gene set, the four immune signatures are also highly consistent in some types of tumors, such as Bladder-TCC, Lung-SCC, and Skin-Melanoma, and EMT may have important roles in the immune microenvironment in these tumors (Hugo et al., 2016; Chae et al., 2018). For the TGFβ signaling gene set, diverse associations with the four immune signatures were detected. WNT/β-catenin signaling was inversely related to these immune signatures and IEI in several tumor types, such as Skin-Melanoma. In Lung-AdenoCA, Lung-SCC, Lymph-BNHL, and Skin-Melanoma, the trends in IEI seemed to be consistent with the four immune signatures.

Further investigations of infiltrated immune cells are important to understand immune escape mechanisms. Based on the predicted composition of infiltrated immune cells and the expression of CD45, a pan-lymphocyte marker, we evaluated the activity of infiltrated immune cells (Supplementary Figure 7). We focused on the activity of M2 macrophages (*y*-axis) as immune suppressive cells and CD8^+^ T-cells (*x*-axis) as immune effector cells, and obtained a flow cytometry-like plot for each tumor type (Figure 4a). We focused on eight types of tumors, Breast-AdenoCA, Cervix-SCC, ColoRect-AdenoCA, Liver-HCC, Lung-AdenoCA, Lung-SCC, Skin-Melanoma, and Uterus-AdenoCA, for finding associations of the immune cell infiltrations to the selective copy number gain and IEI. Lung-AdenoCA and Lung-SCC with selective copy number gains of *TGFB2* (red circles in the upper panels of Figure 4a) showed high activity of M2 macrophages and low activity of CD8^+^ T-cells (statistical significance for repression of CD8^+^ T-cell in selective copy number gain tumors: p = 0.0291 and 0.0244 for the lung AdenoCA and SCC, respectively); in ColoRect-AdenoCA, the selective copy number gain of *TGFB2* was not observed in most samples, and only a small fraction of CD8^+^ T-cell infiltrated tumors with a selective copy number gain of *TGFB2* (p = 0.00453). By contrast, in ColoRect-AdenoCA, infiltrating CD8^+^ T-cells seemed to be repressed in IEI-positive tumors (lower panels of Figure 4a, p = 6.58E-4 for IEI-positive tumors’ CD8+ T-cell repression). In Uterus-AdenoCA, IEI positive tumors also showed a small fraction of CD8+ T-cells (p= 0.00208). These analyses indicated that selective copy number changes and immune escape histories (IEI) of each tumor can reflect the immune cell composition and immune microenvironment within tumors.

**Figure 4:**
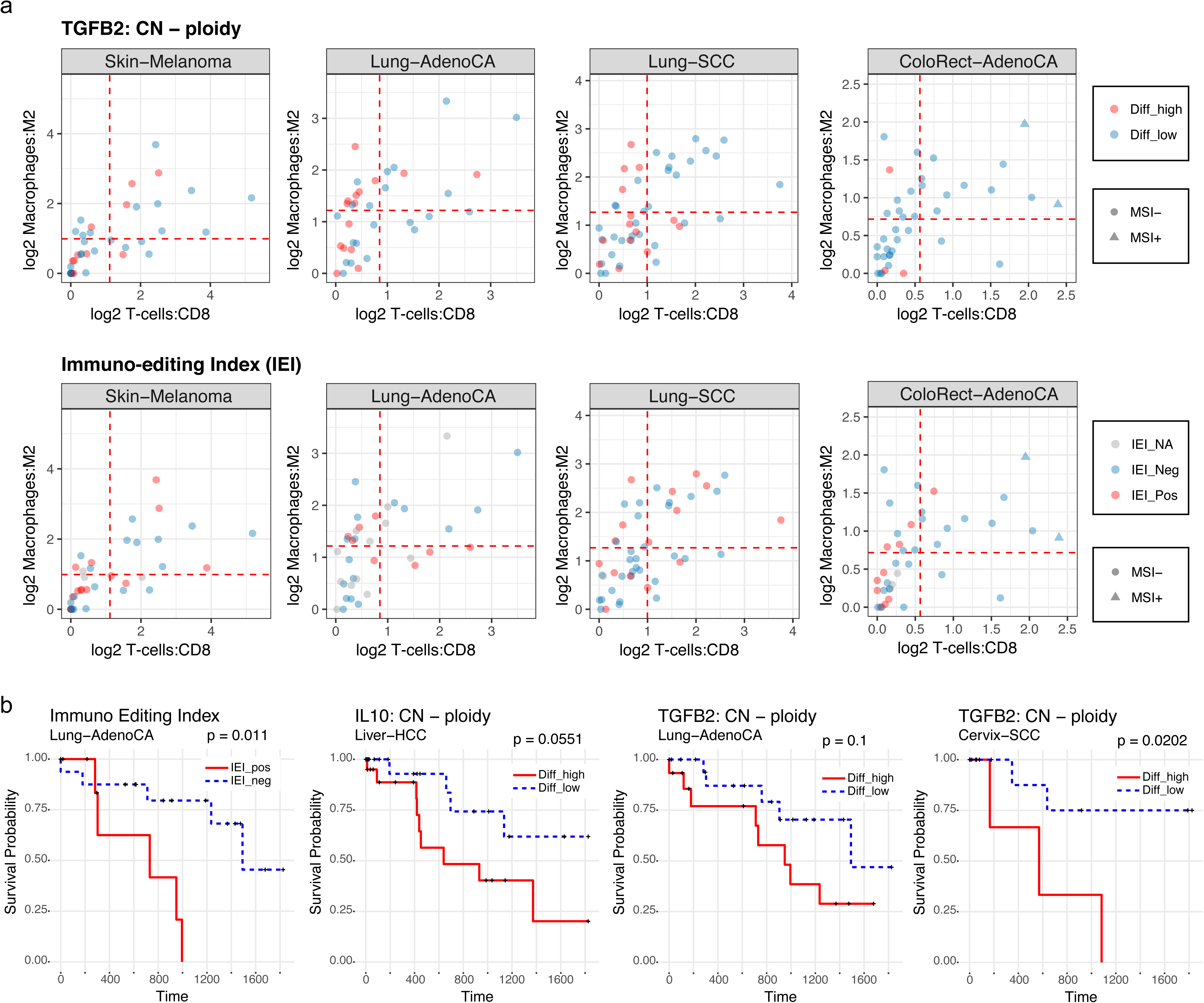
Associations among genomic alterations, immuno-editing history, infiltrated immune cell activities and clinical outcome. (a) Flow cytometry-like plots representing the estimated activity of infiltrated CD8+ T-cells (*x*-axis) and M2 macrophages (*y*-axis). The dotted red line and circle represent the mean value for each axis and a sample, respectively. (b) Kaplan–Meier curves for overall survival show a high IEI and selected copy number gains of *IL10* and *TGFB2*.

**Supplementary Figure 7:**
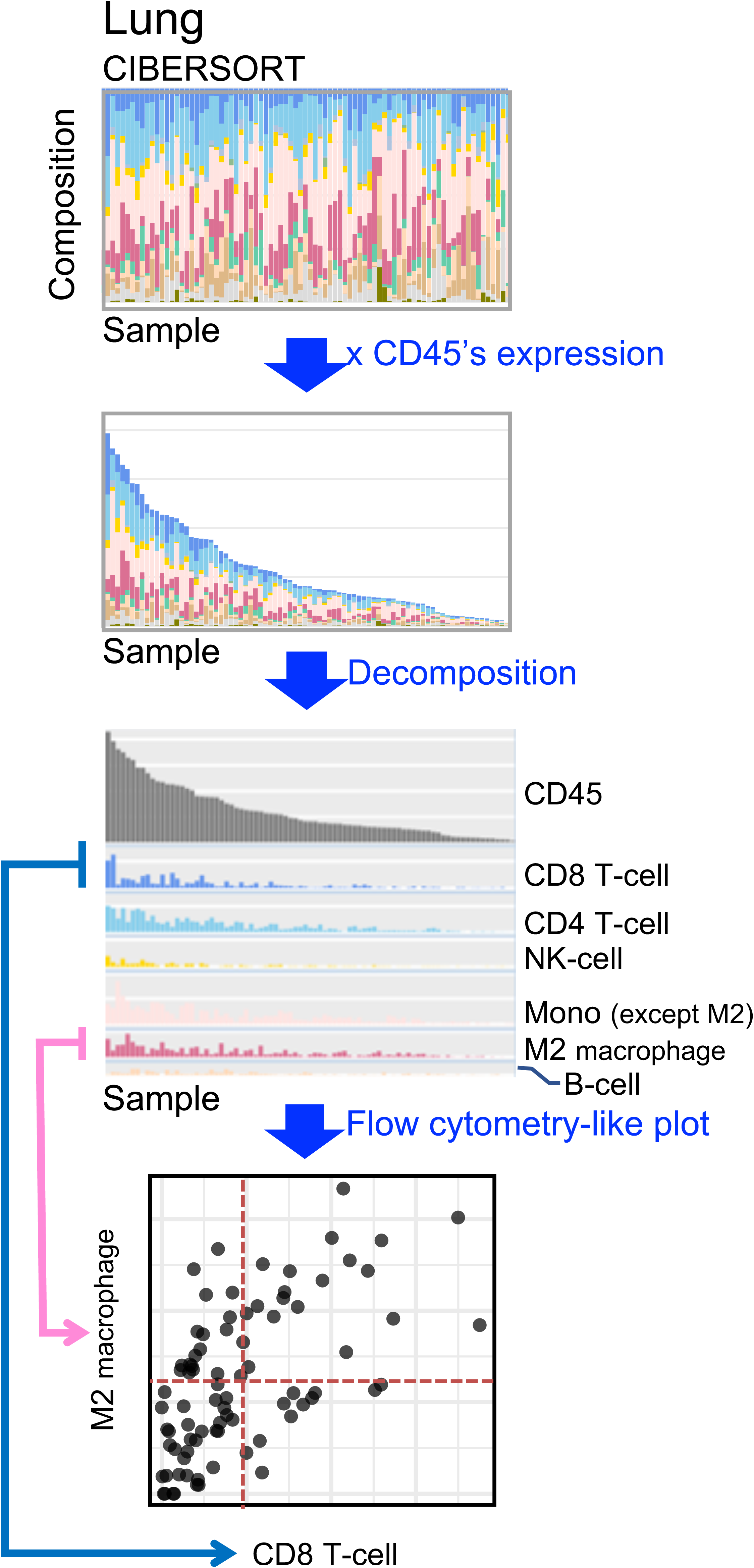
Analysis of infiltrated cells and their predicted activities. Using the results of CIBERSORT and the expression of CD45 for each sample, we estimated the activity of infiltrated immune cells, including CD8+ T-cells, CD4+ T-cells, NK-cells, M2 macrophages, B-cells, etc. An example of a scatter plot (*x*-axis and *y*-axis indicate the predicted activity of CD8+ T-cells and M2 macrophages, respectively) is shown at the bottom, where a circle represents a sample.

**Supplementary Figure 8:**
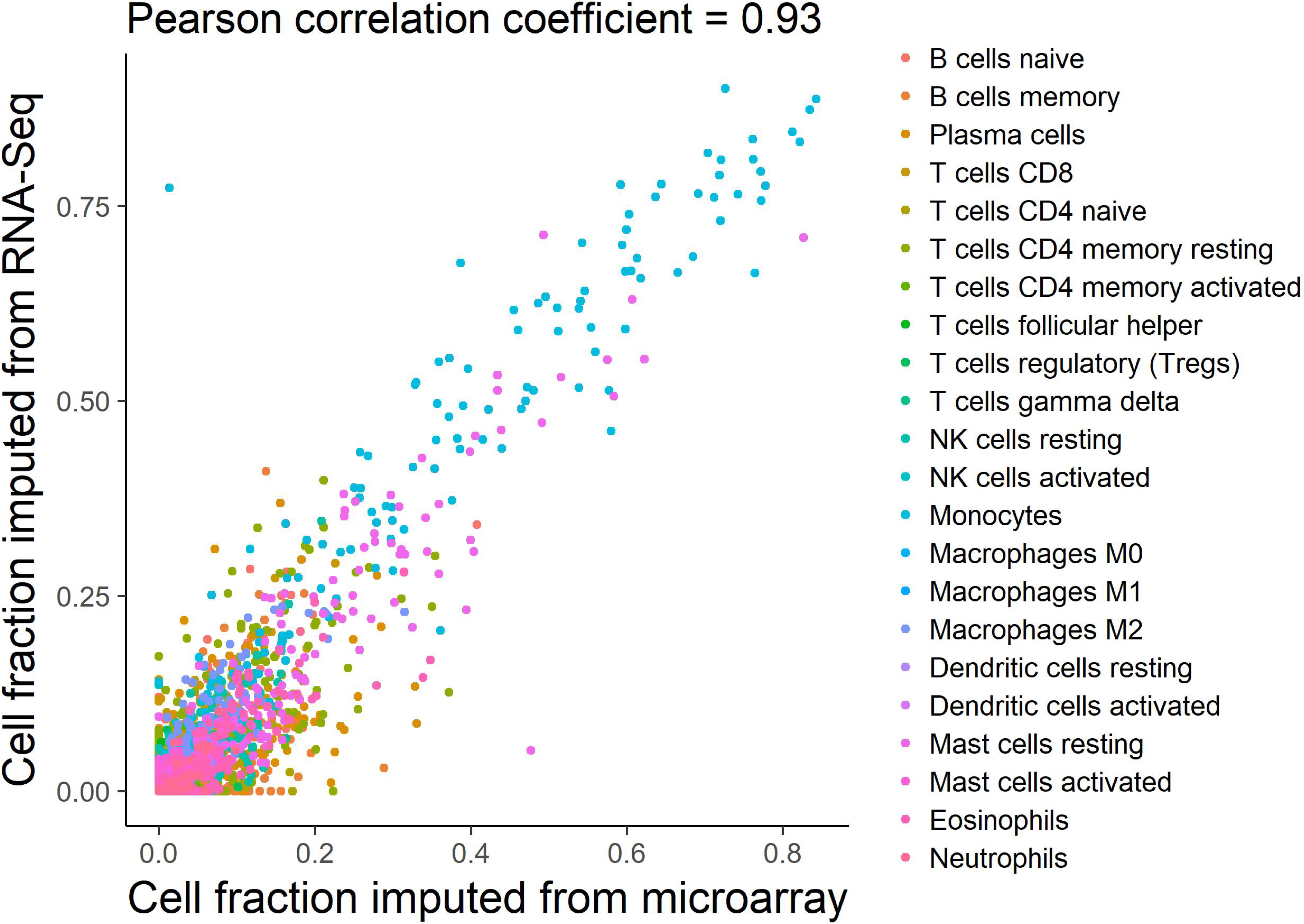
CIBERSORT deconvolution for the comparison between microarray data and RNA-Seq data using 166 TCGA LAML-US samples. Pearson’s correlation coefficients were used to measure concordance.

Finally, we performed a survival analysis of donors partitioned by IEI values for the above eight types of tumors and found that Lung-AdenoCA cancer donors with IEI-positive tumors (immunoediting resistant tumor) exhibited a much worse overall survival than that of donors with IEI-negative tumors. In Lung-AdenoCA, IEI showed a possible separation (p = 0.011, Figure 4b), whereas those for the other aforementioned gene set signatures were not significant. We also analyzed the relationship between selective copy number gain (*IL10* and *TGFB2*) and overall survival and showed three examples using Liver-HCC, Lung-AdenoCA, and Cervix-SCC. For these tumor types, tumors with selective copy number gains of *IL10* or *TGFB2* showed worse overall survival than that of the tumors without these copy number gains (p = 0.0551 (*IL10*) for Liver-HCC, p = 0.1 (*TGFB2*) for Lung-AdenoCA, and p = 0.0202 (*IL10* and *TGFB2*) for Cervix-SCC).

## Conclusion

We derived immuno-genomic profiles, including somatic mutations in immune genes, HLA genotypes, NAGs, and immune micro-environmental landscapes, from pan-cancer whole genome and RNA sequence data. We observed that tumors acquired many types of immune escape mechanisms by selective copy number gains of immune-related genes, failure of the antigen presentation system, and alterations in immune checkpoint molecules in a tumor-specific manner. The history of immunoediting, as estimated using pseudogenes as sites free of immune pressure, indicated associations between tumorigenesis and immune escape across various tumor types. Furthermore, the micro-environmental landscape related to immune characteristics revealed diverse background or intrinsic pathways controlling the non-inflamed subset of each tumor type. This provides essential information for identifying therapeutic targets. These analyses revealed the impact of the immune micro-environment on the immune resistance and/or immune escape of tumors.

## ACKNOWLEDGEMENTS

This study was supported in party by the Japan Agency for Medical Research and Development (AMED) Project for Cancer Research and Therapeutic Evolution (P-CREATE) (to H.N.) and a Grant-in-Aid for Scientific Research (B) (to S.I.) from the JSPS. The super-computing resource ‘SHIROKANE’ was provided by Human Genome Center, The University of Tokyo (http://sc.hgc.jp/shirokane.html).

## METHODS

### Genomic alterations in immune-related genes in pan-cancer datasets

Datasets of somatic point mutations, structural variants (SVs), and copy number alterations were generated as part of the Pan-Cancer Analysis of Whole Genomes (PCAWG) project. Overall, 2834 samples with whole genome data are represented in the PCAWG datasets, spanning a range of cancer types (bladder, sarcoma, breast, liver-biliary, cervix, leukemia, colorectal, lymphoma, prostate, esophagus, stomach, central nervous system, head/neck, kidney, lung, melanoma, ovary, pancreas, thyroid, and uterus). The consensus somatic SVs, CNAs, and SNVs in PCAWG samples were determined by three different data centers using different algorithms; calls made by at least two algorithms were used in downstream analyses. To determine copy number, the calls made by the Sanger group were used (Yang et al., PCAWG Tech paper)

### Hla genotyping and mutations from whole genome sequences

For HLA genotyping using whole genome sequencing data, a Bayesian method known as ALPHLARD (Hayashi et al., 2018) was used; this method was designed to perform accurate HLA genotyping from short-read data and to predict the HLA sequences of the sample. The latter function enables the identification of somatic mutations by comparisons of the HLA sequences of the tumor sample with those of the matched-normal sample. The statistical formulation for the posterior probability can be described as follows:

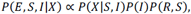

where *R* = (*R*_1_, *R*_2_) is the pair of HLA types (reference sequences), *S* = (*S*_1_, *S*_2_) is the pair of HLA sequences of the samples, *X* = (*x*_1_, *x*_2_, …) is the set of sequence reads, and *I* = (*I*_1_, *I*_2_, …) is the set of variables taking 1 or 2 (the *j*th element, *I_j_* indicates the *j*th read *x_j_* is generated from). On the right-hand side of the above equation, the left term indicates the likelihood of the sequence reads when the HLA sequences and the reference sequences are fixed. The middle and the right terms are the priors. The parameters, HLA sequences, and HLA types, were determined using the MCMC procedure with parallel tempering.

### Immune signatures from RNA-seq data

To investigate the microenvironment related to the immune characteristics of tumors, the following immune-related signatures were prepared (Supplementary Table 2):

- Cytotoxic (Rooney et al., 2015)
- Immune checkpoints (Mahoney et al., 2015; Smyth et al., 2015)
- HLA pathway class I (Neefjes et al., 2011)
- Cell component

Using a signature, two subsets of samples were defined, a subgroup of samples with immune characteristics indicating the focused signature, and a subgroup lacking these characteristics. By comparing RNA expression levels in these subgroups, enriched gene sets or pathways were identified as related microenvironments.

### Immuno-signature-based GSEA

For each cancer type, the samples were divided into two groups based on gene expression patterns of an immuno-signature set, *e.g.*, cytotoxic signature set, and a GSEA was conducted for gene sets using MSigDB by comparing whole gene expression values between the two groups of samples. To obtain the two groups, hierarchical clustering was applied to the gene expression matrix for immunosignature genes of the samples and the dendrogram for the samples was cut at the root. The group with a higher mean expression value for immuno-signature genes than that in the other group was labeled “High,” while the other was labeled “Low.” In this study, as described above, we considered four immuno-signature sets. For an immuno-signature set, the above GSEA was applied for each cancer type and the enrichment results for MSigDB gene sets were compiled into heatmaps. In a heatmap, each cell corresponds to a pair of an MSigDB gene set and cancer type, and has the value of, where the nominal p-value and is an indicator variable; if the gene set is enriched in “High” group and otherwise (**Supplementary Figure 6**).

### Immune cell components

For CIBERSORT implementation, FPKM values were used after upper-quartile normalization as input gene expression values (FPKMs are in linear space, without log-transformation) and the default LM22 was used as the signature gene matrix. Twenty-two leukocyte fractions were imputed from CIBERSORT. Originally, CIBERSORT was proposed for RNA expression data obtained by microarray. However, it has been reported that CIBERSORT can be applied to bulk tumor RNA-seq (Tuong et al., 2016; Mehnert et al., 2016) and single-cell RNA-seq (Baron et al., 2016). The correlation between results obtained using microarray data and RNA-seq data from 166 LAML-US tumors was independently evaluated; the observed correlation coefficient was 0.93, which was significantly high. Therefore, CIBERSORT was applied to RNA-seq data (**Supplementary Figure 8**).

### Neo-antigen prediction

From PCAWG preliminary consensus files, 2,786 annotated .tsv files were generated using ANNOVAR and exclusion samples were removed according to release_may2016.v1.3.tsv. Next, focusing on nonsynonymous mutations in exonic regions, the corresponding mutant/wild-type peptides of length 8–11-mer including an amino-acid substitution were constructed using the UCSC RefSeq mRNA and refFlat data (http://hgdownload.soe.ucsc.edu/downloads.html). Next, binding affinities (IC_50_) of all generated peptides were predicted using netMHCpan3.0 (Nielsen and Andreatta, 2016) for HLA class I and netMHCIIpan3.1 (Andreatta et al., 2015) for HLA class II. Finally, neoantigens were counted for each patient by considering that mutant peptides with IC_50_ values of less than 500 as neoantigens. Here, neoantigens were counted as the number of mutations that can generate neoantigens; thus, each mutation was counted once, even if it generated more than one neoantigen for one or more HLAs. Note that mutations in which annotated information was not consistent with UCSC RefSeq mRNA and refFlat data were skipped as database mismatches. The ratio of the number of non-skipped nonsynonymous mutations to the number of all observed nonsynonymous mutations was defined as the concordance rate. Although this value was nearly 1 in all cases (greater than 0.99, on average), it was used as a tuning parameter, as described below.

### Immuno-editing index

To evaluate the sample-specific immuno-editing history, an immuno-editing index (IEI) describing the degree of accumulated immune suppression was established. IEI compares the ratio of the number of neoantingens to the number of nonsynonymous mutations in exonic regions and in the control regions, which are not affected by immune pressure. Pseudogene regions were used as internal controls for a tumor and only pseudogene mutations whose genomic positions were downstream of the stop codon were extracted according to PseudoPipe v.74 (http://www.pseudogene.org/pseudopipe/). In this concept, the following assumptions were made: (i) nonsynonymous mutations in exonic regions can be suppressed by immune pressure if their mutant peptides can bind to HLAs and (ii) synonymous mutations in exonic regions and nonsynonymous/synonymous mutations in pseudogene regions are not affected by immune pressure. Under these assumptions, the number of nonsynonymous mutations in exonic regions can be lower than the number of ideal nonsynonymous mutations in exonic regions, indicating the hypothetical number of nonsynonymous mutations under non-immune pressure. Several quantities were defined as follows:

- Number of nonsynonymous mutations used to evaluate neoantigens (not skipped by database mismatch) in exonic regions = #nonsynE
- Number of synonymous mutations in exonic regions = #synE
- Number of predicted neoantigens in exonic regions = #NagE
- Number of nonsynonymous mutations used to evaluate neoantigens (not skipped by database mismatch) in pseudogene regions = #nonsynP
- Number of synonymous mutations in pseudogene regions = #synP
- Number of predicted neoantigens in pseudogene regions = #NagP
- Concordance rate of mutation annotations in exonic regions = *c*_exon_
- Concordance rate of mutation annotations in pseudogene regions = *c*_pseudo_

The number of nonsynonymous mutations in exonic region was adjusted to obtain the number of ideal nonsynonymous mutations (#*InonsynE*) using the above quantities as follows:

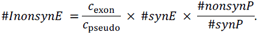

Here, #*InonsynE* was set to #*NagE* if #*InonsynE* was less than #*NagE*. IEI was calculated as the modified log ratio in terms of the numbers of neoantingens and nonsynonymous mutations, and is equal to the sum of the numbers of neoantingens and nonneoantingens between exonic and pseudogene regions as follows:

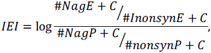

where *C* is a regularized constant, set to 0.5 for the analysis.

### Pseudogene selection

PseudoPipe (build 74) (Zhang et al., 2006) was used as a pseudogene database for the following analysis, which includes the region and the parental gene of each pseudogene, among other information. First, pseudogene mutations in each sample were extracted from the VCF file based on pseudogene regions described in PseudoPipe. Next, each pseudogene in PseudoPipe was aligned to the parental gene using Clustal Omega (version 1.2.1) (Sievers et al., 2011) with default settings. Each pseudogene mutation was converted to a parental gene mutation located at the same position as that of the pseudogene mutation in the alignment. Note that pseudogene mutations were excluded in the following neo-antigen analysis if the position corresponded to an intron of the parental gene or if the bases differed at the position in the alignment of the pseudogene and the parental gene. Thus, except for the above cases, pseudogene mutations were treated as if they were exonic mutations. An immuno-editing history analysis was applied to the converted mutations and the results were used as an internal control. Mutations in pseudogene regions were used directly, without information for parental genes. However, the amino acid composition in pseudogene regions with parental genes is considered similar to that in exonic regions. Additionally, in pseudogene regions, many stop codons are present and a method was determined to handle these. Therefore, pseudogene regions with parental genes were used as a suitable internal control to evaluate the strength of immune pressure.

## REFERENCES

1. Hanahan D, Weinberg RA. (2011) Hallmarks of cancer: The next generation. Cell 144(5), 646–674.

2. Lefranc MP, Giudicelli V, Ginestoux C, Bodmer J, Müller W, Bontrop R, Lemaitre M, Malik A, Barbié V, Chaume D. (1999) IMGT, the international ImMunoGeneTics database. Nucleic Acids Res 27, 209–212.

3. Linnemann C, van Buuren MM, Bies L, Verdegaal EM, Schotte R, Calis JJ, Behjati S, Velds A, Hilkmann H, Atmioui DE, Visser M, Stratton MR, Haanen JB, Spits H, van der Burg SH, Schumacher TN. (2015) High-throughput epitope discovery reveals frequent recognition of neo-antigens by CD4+ T cells in human melanoma. Nat Med 21, 81–85.

4. Tran E, Turcotte S, Gros A, Robbins PF, Lu YC, Dudley ME, Wunderlich JR, Somerville RP, Hogan K, Hinrichs CS, Parkhurst MR, Yang JC, Rosenberg SA. (2014) Cancer immunotherapy based on mutation-specific CD4+ T cells in a patient with epithelial cancer. Science 344, 641–645.

5. Kreiter S, Vormehr M, van de Roemer N, Diken M, Löwer M, Diekmann J, Boegel S, Schrörs B, Vascotto F, Castle JC, Tadmor AD, Schoenberger SP, Huber C, Türeci Ö, Sahin U. (2015) Mutant MHC class II epitopes drive therapeutic immune responses to cancer. Nature 520, 692–696.

6. Yarchoan M, Johnson BA 3rd, Lutz ER, Laheru DA, Jaffee EM. (2017) Targeting neoantigens to augment antitumour immunity. Nat Rev Cancer 17, 209–222.

7. Grivennikov SI, Greten FR, Karin M. (2010) Immunity, inflammation, and cancer. Cell 140(6):883–899.

8. Schreiber RD, Old LJ, Smyth MJ. (2011) Cancer immunoediting: integrating immunity’s roles in cancer suppression and promotion. Science 331(6024), 1565–1570

9. Sharma P, Wagner K, Wolchok JD, Allison JP. (2011) Novel cancer immunotherapy agents with survival benefit: recent successes and next steps. Nat Rev Cancer 11, 805–812.

10. Pardoll DM. (2012) The blockade of immune checkpoints in cancer immunotherapy. Nat Rev Cancer 12, 252–264.

11. Mahoney KM, Rennert PD, Freeman GJ. (2015) Combination cancer immunotherapy and new immunomodulatory targets. Nat Rev Drug Discov, 14, 561–584.

12. Zaretsky JM, Garcia-Diaz A, Shin DS, Escuin-Ordinas H, Hugo W, Hu-Lieskovan S, Torrejon DY, Abril-Rodriguez G, Sandoval S, Barthly L, Saco J, Homet Moreno B, Mezzadra R, Chmielowski B, Ruchalski K, Shintaku IP, Sanchez PJ, Puig-Saus C, Cherry G, Seja E, Kong X, Pang J, Berent-Maoz B, Comin-Anduix B, Graeber TG, Tumeh PC, Schumacher TN, Lo RS, Ribas A. (2016) Mutations associated with acquired resistance to PD-1 blockade in melanoma. N Engl J Med, 375, 819–829.

13. Anagnostou V, Smith KN, Forde PM, Niknafs N, Bhattacharya R, White J, Zhang T, Adleff V, Phallen J, Wali N, Hruban C, Guthrie VB, Rodgers K, Naidoo J, Kang H, Sharfman W, Georgiades C, Verde F, Illei P, Li QK, Gabrielson E, Brock MV, Zahnow CA, Baylin SB, Scharpf RB, Brahmer JR, Karchin R, Pardoll DM, Velculescu VE. (2017) Evolution of neoantigen landscape during immune checkpoint blockade in non-small cell lung cancer. Cancer Discov, 7, 264–276.

14. Gao J, Shi LZ, Zhao H, Chen J, Xiong L, He Q, Chen T, Roszik J, Bernatchez C, Woodman SE, Chen PL, Hwu P, Allison JP, Futreal A, Wargo JA, Sharma P. (2016) Loss of IFN-γ pathway genes in tumor cells as a mechanism of resistance to anti-CTLA-4 therapy. Cell 167, 397–404.

15. Shin DS, Zaretsky JM, Escuin-Ordinas H, Garcia-Diaz A, Hu-Lieskovan S, Kalbasi A, Grasso CS, Hugo W, Sandoval S, Torrejon DY, Palaskas N, Rodriguez GA, Parisi G, Azhdam A, Chmielowski B, Cherry G, Seja E, Berent-Maoz B, Shintaku IP, Le DT, Pardoll DM, Diaz LA Jr, Tumeh PC, Graeber TG, Lo RS, Comin-Anduix B, Ribas A. (2017) Primary resistance to PD-1 blockade mediated by JAK1/2 mutations. Cancer Discov 7, 188–201.

16. Davoli T, Uno H, Wooten EC, Elledge SJ. (2017) Tumor aneuploidy correlates with markers of immune evasion and with reduced response to immunotherapy. Science 355(6322), eaaf8399.

17. Itakura E, Huang RR, Wen DR, Paul E, Wünsch PH, Cochran AJ. (2011) IL-10 expression by primary tumor cells correlates with melanoma progression from radial to vertical growth phase and development of metastatic competence. Mod Pathol, 24, 801–809.

18. Wiguna AP and Walden P. (2015) Role of IL-10 and TGF-β in melanoma. Exp Dermatol, 24, 209–214.

19. Yang L, Pang Y, Moses HL. (2015) TGF-beta and immune cells: an important regulatory axis in the tumor microenvironment and progression. Trends Immunol, 31, 220–227.

20. Lan Y, Zhang D, Xu C, Hance KW, Marelli B, Qi J, Yu H, Qin G, Sircar A, Hernández VM, Jenkins MH, Fontana RE, Deshpande A, Locke G, Sabzevari H, Radvanyi L, Lo KM. (2018) Enhanced preclinical antitumor activity of M7824, a bifunctional fusion protein simultaneously targeting PD-L1 and TGF-β. Sci Transl Med, 10, pii: eaan5488.

21. Strauss J, Heery CR, Schlom J, Madan RA, Cao L, Kang Z, Lamping E, Marté JL, Donahue RN, Grenga I, Cordes L, Christensen O, Mahnke L, Helwig C, Gulley JL. (2018) Phase I trial of M7824 (MSB0011359C), a bifunctional fusion protein targeting PD-L1 and TGFβ, in advanced solid tumors. Clin Cancer Res, 24, 1287–1295.

22. Campbell P, et al., PCAWG marker paper. in submission.

23. Zhang Y, Chen F, Fonseca NA, He Y, Fujita M, Nakagawa H, Zhang Z, Brazma A, Creighton C. Whole genome and RNA sequencing of 1, 220 cancers reveals hundreds of genes deregulated by rearrangement of cis-regulatory elements, PCAWG marker paper. Biorxiv, doi: https://doi.org/10.1101/099861

24. Hackl H, Charoentong P, Finotello F, Trajanoski Z. (2016) Computational genomics tools for dissecting tumour-immune cell interactions. Nat Rev Genet 17(8):441–458.

25. Laumont CM, Vincent K, Hesnard L, Audemard É, Bonneil É, Laverdure JP, Gendron P, Courcelles M, Hardy MP, Côté C, Durette C, St-Pierre C, Benhammadi M, Lanoix J, Vobecky S, Haddad E, Lemieux S, Thibault P, Perreault C. (2018) Noncoding regions are the main source of targetable tumor-specific antigens. Sci Transl Med. Dec 5;10(470). pii: eaau5516. doi: 10.1126/scitranslmed.aau5516.

26. Kataoka K, Shiraishi Y, Takeda Y, Sakata S, Matsumoto M, Nagano S, Maeda T, Nagata Y, Kitanaka A, Mizuno S, Tanaka H, Chiba K, Ito S, Watatani Y, Kakiuchi N, Suzuki H, Yoshizato T, Yoshida K, Sanada M, Itonaga H, Imaizumi Y, Totoki Y, Munakata W, Nakamura H, Hama N, Shide K, Kubuki Y, Hidaka T, Kameda T, Masuda K, Minato N, Kashiwase K, Izutsu K, Takaori-Kondo A, Miyazaki Y, Takahashi S, Shibata T, Kawamoto H, Akatsuka Y, Shimoda K, Takeuchi K, Seya T, Miyano S, and Ogawa S. (2016) Aberrant PD-L1 expression through 3?-UTR disruption in multiple cancers. Nature 534(7607), 402–406.

27. Latchman Y, Wood CR, Chernova T, Chaudhary D, Borde M, Chernova I, Iwai Y, Long AJ, Brown JA, Nunes R, Greenfield EA, Bourque K, Boussiotis VA, Carter LL, Carreno BM, Malenkovich N, Nishimura H, Okazaki T, Honjo T, Sharpe AH, Freeman GJ. (2001) PD-L2 is a second ligand for PD-1 and inhibits T cell activation. Nat Immunol 2(3), 261–268.

28. Rozali EN, Hato SV, Robinson BW, Lake RA, Lesterhuis WJ. (2012) Programmed death ligand 2 in cancer-induced immune suppression. Clin Dev 2012, 656340.

29. Jahnke M, Trowsdale J, Kelly AP (2012). Structural requirements for recognition of major histocompatibility complex class II by membrane-associated ring-ch (march) protein e3 ligases. J Biol Chem 287(34), 28779–28789.

30. Albring J, Koopmann JO, Hämmerling GJ, Momburg F. (2004) Retrotranslocation of mhc class i heavy chain from the endoplasmic reticulum to the cytosol is dependent on ATP supply to the ER lumen. Mol Immunol 40(10), 733–741.

31. Le DT, Uram JN, Wang H, Bartlett BR, Kemberling H, Eyring AD, Skora AD, Luber BS, Azad NS, Laheru D, Biedrzycki B, Donehower RC, Zaheer A, Fisher GA, Crocenzi TS, Lee JJ, Duffy SM, Goldberg RM, de la Chapelle A, Koshiji M, Bhaijee F, Huebner T, Hruban RH, Wood LD, Cuka N, Pardoll DM, Papadopoulos N, Kinzler KW, Zhou S, Cornish TC, Taube JM, Anders RA, Eshleman JR, Vogelstein B, Diaz LA Jr. (2015) PD-1 blockade in tumors with mismatch-repair deficiency. N Engl J Med 372(26), 2509–2520.

32. Fujimoto A, et al., (2017) PCAWG marker paper. in preparation.

33. Robbins PF, Lu YC, El-Gamil M, Li YF, Gross C, Gartner J, Lin JC, Teer JK, Cliften P, Tycksen E, Samuels Y, Rosenberg SA. (2013) Mining exomic sequencing data to identify mutated antigens recognized by adoptively transferred tumor-reactive T cells. Nat Med 19(6), 747–752.

34. Carreno BM, Magrini V, Becker-Hapak M, Kaabinejadian S, Hundal J, Petti AA, Ly A, Lie WR, Hildebrand WH, Mardis ER, Linette GP. (2015) Cancer immunotherapy. A dendritic cell vaccine increases the breadth and diversity of melanoma neoantigen-specific T cells. Science 348(6236), 803–808.

35. Burnet FM. (1970) The concept of immunological surveillance. Prog Exp Tumor Res 13, 1–27.

36. Dunn GP, Bruce AT, Ikeda H, Old LJ, Schreiber RD. (2002) Cancer immunoediting: from immunosurveillance to tumor escape. Nat Immunology 3(11), 991–998.

37. Terry S, Savagner P, Ortiz-Cuaran S, Mahjoubi L, Saintigny P, Thiery JP, Chouaib S. New insights into the role of EMT in tumor immune escape. Mol Oncol, 11, 824–846.

38. Pai SG, Carneiro BA, Mota JM, Costa R, Leite CA, Barroso-Sousa R, Kaplan JB, Chae YK, Giles FJ. (2017) Wnt/beta-catenin pathway: modulating anticancer immune response. J Hematol Oncol, 10, 101.

39. Hugo W, Zaretsky JM, Sun L, Song C, Moreno BH, Hu-Lieskovan S, Berent-Maoz B, Pang J, Chmielowski B, Cherry G, Seja E, Lomeli S, Kong X, Kelley MC, Sosman JA, Johnson DB, Ribas A, Lo RS. (2016) Genomic and transcriptomic features of response to anti-PD-1 therapy in metastatic melanoma. Cell 165, 35–44.

40. Chae YK, Chang S, Ko T, Anker J, Agte S, Iams W, Choi WM, Lee K, Cruz M. (2018) Epithelial-mesenchymal transition (EMT) signature is inversely associated with T-cell infiltration in non-small cell lung cancer (NSCLC). Sci Rep 8, 2918.

41. Spranger S, Bao R, Gajewski TF. (2015) Melanoma-intrinsic β-catenin signalling prevents anti-tumour immunity. Nature 523, 231–235.

42. Newman AM, Liu CL, Green MR, Gentles AJ, Feng W, Xu Y, Hoang CD, Diehn M, Alizadeh AA. (2015) Robust enumeration of cell subsets from tissue expression profiles. Nat Methods 12(5), 453–457.

## REFERENCES in METHOD SECTION

1. Hayashi S, Yamaguchi R, Mizuno S, Komura M, Miyano S, Nakagawa H, Imoto S. (2018) ALPHLARD: a Bayesian method for analyzing HLA genes from whole genome sequence data, BMC Genomics, 19(1):790.

2. Rooney MS, Shukla SA, Wu CJ, Getz G, Hacohen N. (2015) Molecular and genetic properties of tumors associated with local immune cytolytic activity. Cell, 160, 48–61.

3. Smyth MJ, Ngiow SF, Ribas A, Teng MW. (2015) Combination cancer immunotherapies tailored to the tumour microenvironment. Nat Rev Clin Oncol, 13, 143–158.

4. Neefjes J, Jongsma ML, Paul P, Bakke O. (2011) Towards a systems understanding of MHC class I and MHC class II antigen presentation. Nat Rev Immunol, 11, 823–836.

5. Tuong ZK, Fitzsimmons R, Wang SM, Oh TG, Lau P, Steyn F, Thomas G, Muscat GE. (2016) Transgenic adipose-specific expression of the nuclear receptor RORα drives a striking shift in fat distribution and impairs glycemic control. EbioMedicine, 11, 101–117.

6. Mehnert JM, Panda A, Zhong H, Hirshfield K, Damare S, Lane K, Sokol L, Stein MN, Rodriguez-Rodriquez L, Kaufman HL, Ali S, Ross JS, Pavlick DC, Bhanot G, White EP, DiPaola RS, Lovell A, Cheng J, Ganesan S. (2016) Immune activation and response to pembrolizumab in POLE-mutant endometrial cancer. J Clin Invest, 126, 2334–23140.

7. Baron M, Veres A, Wolock SL, Faust AL, Gaujoux R, Vetere A, Ryu JH, Wagner BK, Shen-Orr SS, Klein AM, Melton DA, Yanai I. (2016) A single-cell transcriptomic map of the human and mouse pancreas reveals inter‐ and intra-cell population structure, Cell Syst, 3, 346–360.e4.

8. Nielsen M, Andreatta M. (2015) NetMHCpan-3.0; improved prediction of binding to MHC class I molecules integrating information from multiple receptor and peptide length datasets. Genome Med, 8, 33.

9. Andreatta M, Karosiene E, Rasmussen M, Stryhn A, Buus S, and Nielsen M. (2015) Accurate pan-specific prediction of peptide-MHC class II binding affinity with improved binding core identification. Immunogenetics, 67, 641–650.

10. Zhang Z, Carriero N, Zheng D, Karro J, Harrison PM, Gerstein M. (2006) PseudoPipe: an automated pseudogene identification pipeline. Bioinformatics, 22, 1437–1439.

11. Sievers F, Wilm A, Dineen D, Gibson TJ, Karplus K, Li W, Lopez R, McWilliam H, Remmert M, Söding J, Thompson JD, Higgins DG. (2011) Fast, scalable generation of high-quality protein multiple sequence alignments using Clustal Omega. Mol Syst Biol, 7, 539.

